# Joint species distribution modelling with HMSC-R

**DOI:** 10.1101/603217

**Authors:** Gleb Tikhonov, Øystein Opedal, Nerea Abrego, Aleksi Lehikoinen, Otso Ovaskainen

**Affiliations:** Department of Computer Science, Aalto University, FI-02150, Espoo, Finland; Organismal and Evolutionary Biology Research Programme, PO Box, 65, FI-00014 University of Helsinki, Finland; Centre for Biodiversity Dynamics, Department of Biology, Norwegian University of Science and Technology, N-7491 Trondheim, Norway; Department of Agricultural Sciences, PO Box 27, FI-00014 University of Helsinki, Finland; The Helsinki Lab of Ornithology, Finnish Museum of Natural History, PO Box 17, FI-00014 University of Helsinki, Finland

**Keywords:** Community modelling, Community similarity, Multivariate data, Species Distribution Modelling

## Abstract

- Joint Species Distribution Modelling (JSDM) is becoming an increasingly popular statistical method for analyzing data in community ecology. JSDM allow the integration of community ecology data with data on environmental covariates, species traits, phylogenetic relationships, and the spatio-temporal context of the study, providing predictive insights into community assembly processes from non-manipulative observational data of species communities. Hierarchical Modelling of Species Communities (HMSC) is a general and flexible framework for fitting JSDMs, yet its full range of functionality has remained restricted to Matlab users only.
- To make HMSC accessible to the wider community of ecologists, we introduce HMSC-R 3.0, a user-friendly R implementation of the framework described in Ovaskainen et al (Ecology Letters, 20 (5), 561-576, 2017) and further extended in several later publications.
- We illustrate the use of the package by providing a series of five vignettes that apply HMSC-R 3.0 to simulated and real data. HMSC-R applications to simulated data involve single-species models, models of small communities, and models of large species communities. They demonstrate the estimation of species responses to environmental covariates and how these depend on species traits, as well as the estimation of residual species associations. They further demonstrate how HMSC-R can be applied to normally distributed data, count data, and presence-absence data. The real data consist of bird counts in a spatio-temporally structured dataset, environmental covariates, species traits and phylogenetic relationships. The vignettes demonstrate how to construct and fit many kinds of models, how to examine MCMC convergence, how to examine the explanatory and predictive powers of the models, how to assess parameter estimates, and how to make predictions.
- The package, along with the extended vignettes, makes JSDM fitting and post-processing easily accessible to ecologists familiar with R.

## Introduction

Assembly processes make ecological communities vary spatio-temporally in the number, abundance, identities and traits of their constituent species (Leibold et al. 2004; Götzenberger et al. 2012; Vellend 2016). Thus, one of the main goals of modern community ecology is to identify and disentangle the assembly processes that contribute to the observed variation among the studied communities. For reaching this objective, model-based probabilistic approaches, and particularly species distribution models (SDMs), are becoming increasingly popular (D’Amen et al. 2017). There are two main SDM-based approaches for analyzing community data, namely stacked species distribution models (SSDMs), and joint species distribution models (JSDMs). SSDMs model each species separately and then combine (‘stack’) their predictions to assess community-level patterns (Guisan & Rahbek 2011; Calabrese et al. 2014), whereas JSDMs explicitly acknowledge the multivariate nature of communities by assuming that the species respond jointly to the environment and to each other (Warton et al. 2015; Clark et al. 2017; Ovaskainen et al. 2017a).

The Hierarchical Modelling of Species Communities (HMSC) framework belongs to the class of JSDMs, and can be used to interrelate data on species occurrence, environmental covariates, species traits and phylogenetic relationships with community assembly processes, and it is applicable to a wide range of common ecological data types (Ovaskainen et al. 2017a). Within HMSC, environmental filtering is modelled at the species level by measuring how the occurrences of each species depends on environmental conditions. These species-level models are incorporated with a hierarchical structure aimed at determining to what extent environmental filtering is structured by species-specific traits, and/or whether phylogenetically related species exhibit shared environmental responses (Abrego, Norberg & Ovaskainen 2017). Biotic interactions and influences of missing environmental covariates are captured by residual species-to-species association matrices, which may be estimated at multiple spatio-temporal scales (Ovaskainen et al. 2016a; Ovaskainen et al. 2016b; Ovaskainen et al. 2017b), as well as for different environmental conditions (Tikhonov et al. 2017).

The HMSC framework handles data that are spatially explicit (Ovaskainen et al. 2016b), spatially hierarchical (Ovaskainen et al. 2016a), temporal (Sebastián-González et al. 2010) or spatio-temporal (Ovaskainen et al. 2017b). Likewise, HMSC handles different response variable types, such as presence-absence data (Ovaskainen et al. 2016a), count data (Häkkilä et al. 2018), and cover data (Tikhonov et al. 2017). Below, we illustrate the R implementation of HMSC using spatially explicit bird count data as an example. Compared to HMSC-R 2.0 (see Ovaskainen et al. 2017a), the current HMSC-R 3.0 version implements several extensions, enabling one to ask how environmental conditions influence species-to-species association matrices (Tikhonov et al. 2017) and to infer species-to-species associations from time-series data of species-rich communities (Ovaskainen et al. 2017b). Furthermore, compared to HMSC-R 2.0, the current HMSC-R 3.0 version offers much improved flexibility with respect to the random error structures, model fitting efficiency, and greater functionality for post-processing the results and for making predictions. To make this possible, HMSC-R 3.0 has been re-coded anew rather than upgraded from HMSC-R 2.0, and its syntax is not compatible to HSMC-R 2.0. To make JSDM parameterization, validation and post-processing of results with HMSC more easily accessible to ecologists familiar with R, the new package comes with five tutorials in vignette format.

## HMSC-R workflow

Running a typical HSMC analysis includes five main steps: (1) Setting model structure and fitting the model, (2) Examining MCMC convergence, (3) Evaluating model fit, (4) Exploring parameter estimates, and (5) Making predictions. Below, we explain each step in turn.

### Step 1. Setting model structure and fitting the model

In this step, the user loads the data and makes decisions about model structure, including random effects, environmental covariates, and the inclusion or exclusion of species traits and phylogenetic relationships. Model fitting includes running the Markov chain Monte Carlo (MCMC) estimation scheme to sample from the posterior distribution of model parameters.

### Step 2. Examining MCMC convergence

In this step, the user examines whether the MCMC scheme has resulted in a valid approximation of the posterior distribution, e.g. in the sense of the chains having reached a stationary distribution and representing a sufficiently large number of effective number of samples. If not, the results will not be reliable, and thus the user should refit the model with a longer MCMC sampling scheme.

### Step 3. Evaluating model fit

HMSC-R comes with built-in functions that can be used to examine different aspects of model fit. Model fit can be evaluated either in terms of explanatory power, i.e. for the same sampling units that were used to fit the model, or in terms of predictive power through cross-validation, i.e. for other sampling units than those used to fit the model. If explanatory power is much higher than predictive power, the specified model is probably too flexible, and the user may wish to re-consider the model structure and/or the types of environmental variables included.

### Step 4. Exploring parameter estimates

In this step, the user can extract numerical summaries of parameter estimates, e.g. posterior means and quantiles. HMSC-R also comes with functions for producing plots that illustrate the posterior distributions of high-dimensional variables, such as variance partitioning of the explained variation among environmental covariates and random effects, the responses of the species to environmental covariates, and species-to-species associations.

### Step 5. Making predictions

HMSC-R comes with a generic predict function as well as more specific tools for generating predictions over environmental gradients. The user can visualize the predictions both at the species level (e.g. occurrence probabilities or abundances of individual species) or at the community level (e.g. species richness or community-weighted trait means). Similarly, predictions over space can be used to generate maps for e.g. distributions of individual species, community weighted trait means, or regions of common profile. Predictions for focal species can also be made conditional on the known occurrences of other species.

We illustrate HMSC-R with simulated data examples in the vignettes, showing how the type and amount of data influences the mixing properties of the MCMC sampling scheme as well as the ability to recover the true parameter values. Below, we illustrate HMSC-R using data on Finnish bird communities.

## Real data example: Finnish birds

### Step 1. Setting model structure and fitting the model

We fitted HMSC to count data on the 50 most common Finnish birds surveyed during 914 counts on 200 permanent transect routes during the years 2006-2014 (same routes counted multiple times; for methodology see Lindström et al. 2015). As environmental covariates (the matrix **X**) we included the categorical variable “habitat type” with the five levels of broadleaved forests (including mixed forest), coniferous forests, open habitats (mountains and scrubland), urban habitats (human settlements and farmland) and wetlands (marine and inland water ecosystems and peatlands), and the continuous covariate “spring temperature” (mean in April and May from European Agency of Climate; Haylock et al. 2008), for which we included also a squared term to allow for intermediate niche optimum. Habitat data was based on Corine land cover data from years 2006 (used for study years 2006–2009) and 2012 (used for study years 2010–2014) (European Environment Agency 2016) and measured within 300 meters buffer from the census sites. As species traits (the matrix **T**), we included the categorical variable “migration strategy” with three levels (resident, short-distance migrant and long-distance migrant, see Saurola et al. 2013, Valkama et al. 2014), and the continuous variable “body size” (log-transformed, according to Cramp et al. 1977–1994). We included in the analyses a phylogenetic tree for the study species, acquired from birdtree.org (Jetz et al. 2012). As community-level random effect, we included the survey route, which we assumed to be spatially structured and hence implemented with the help of spatial latent factors (Ovaskainen et al. 2016b). We fitted both a probit model to data truncated to presence-absence (Model PA), as well as a lognormal Poisson model for the full count data including zeros and non-zeros (Model ABU). For both model types, we considered three model variants that included either both environmental covariates and spatial latent variables (XS), only environmental covariates (X), or only spatial latent variables (S). Thus, we fitted in total the six models PA.XS, PA.X, PA.S, ABU.XS, ABU.X, and ABU.S. We applied the default priors in HMSC, and sampled the posterior distribution with four MCMC chains, each of which were run for 150,000 iterations, out of which the first 50,000 were removed as burn-in and the remaining ones were thinned by 100 to yield 1000 posterior samples per chain, and thus 4000 posterior samples in total. We note that as running long MCMC chains can take a long time, the user is recommended to first test the setting up of the model and explore initial results with shorter runs. For this reason, before fitting the final models, we fitted otherwise identical models with 150 iterations (burn-in 50, thin 1), 1500 iterations (burn-in 500, thin 1) and 15,000 iterations (burn-in 5000, thin 10).

### Step 2. Examining MCMC convergence

The mixing of the MCMC chains was satisfactory but not ideal (Fig. 1AB): the effective sample sizes were much below the theoretical optimum of 4000 samples, and for some parameters the potential scale reduction factor was above 1.1 (see Vignettes). This is because posterior sampling of non-normal models with MCMC for large and multidimensional data is highly challenging in general (Duan et al. 2017), and one key area where further methods development in probabilistic modelling are required. In contrast, the mixing properties of HMSC-R are much better for normally distributed data, as shown by the simulated data examples in the vignettes.

**Figure 1.**
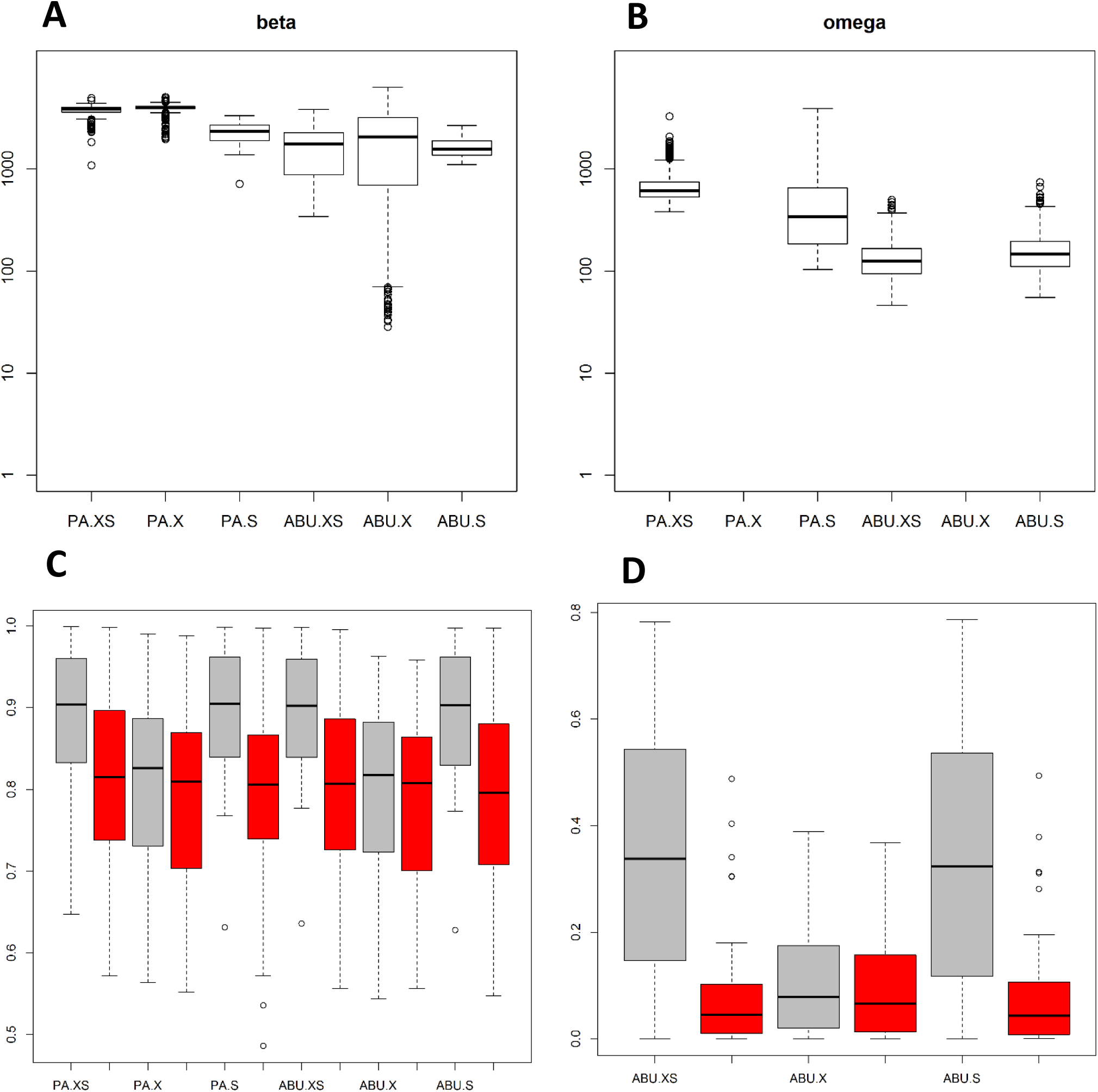
Examining MCMC convergence and evaluation of model fit. The upper panels show the distributions of effective number of posterior samples for two selected multivariate parameters: following the notation of Ovaskainen et al. (2017a), the beta (*β*) parameters (A) model species niches, whereas the omega (Ω) parameters model (B) residual species associations. The lower panels show explanatory (grey bars) and predictive (red bars) power. Panel C shows the ability of all six models to predict species presence-absence in terms of the distributions of species-specific AUC values. Panel D shows for the three abundance models the ability to predict variation in species abundance conditional on presence, measured as squared Spearman rank correlation between prediction and data.

### Step 3. Evaluating model fit

As expected, the explanatory power of the models as evaluated by the AUC statistic (Fig. 1C, grey bars) are always higher than the predictive power (red bars), the latter of which are here based on four-fold cross-validation. We chose four-fold cross-validation as e.g. two-fold cross-validation would include only half of the data for fitting, and thus potentially lead to compromised results, whereas e.g. leave-one-out cross-validation would be computationally very demanding. While the models including spatial components clearly had the highest explanatory power, their predictive power was approximately equal to that of the models that included only environmental covariates, indicating less overfitting of the latter. The lognormal Poisson models were almost equally good in separating presences from absences as the probit models, even while the latter were specifically tailored for doing so. The spatial lognormal Poisson models had relatively high explanatory power, but not predictive power, related to abundance variation (Fig. 1D).

### Step 4. Exploring parameter estimates

While model fit describes how much of the variation in the data the model is able to explain or predict, variance partitioning describes which components of the model that explain the explained variance. For example, in the model PA.XS, the partitioning of the explained variance attributed on average (over the species) 88% to the spatial random effect of the route, and only 5% to climatic and 7% to habitat variables. In contrast, in the model PA.X, that did not include spatial random effects, 72% of the explained variance was attributed to climatic and 28% to habitat variables. Environmental filtering can be assessed by examining the *β*-parameters (regression slopes) that characterize species niches, i.e. the influence of environmental variation on species occurrences (Ovaskainen et al. 2017a). In the model PA.X, many species exhibited statistically well supported responses to many of the covariates, for example a generally negative response to the squared effect of spring temperature (Fig. 2A), suggesting an intermediate optimum. We may next ask how much of the variation in species occurrences can be related to their traits and phylogenetic relationships. In the model PA.X, the included traits explained 7% of the variation in species occurrence, and the residual variation showed no evidence for a phylogenetic signal, as the estimate (posterior median and 95% credibility interval) of the phylogenetic signal parameter *ρ* was 0 (0.00 0.22). The data exhibited a clear spatial signal, as e.g. in the model PA.S the spatial scale parameter *α* related to the leading latent variable was 400 km (200 km… 1200 km). The spatial latent variables also indicated a strong co-occurrence pattern, with a large number of species exhibiting positive associations beyond those explained by the covariates (Fig. 2B). Here the exception was *Phoenicurus phoenicurus*, which was typically recorded on routes where many other species were not recorded. This species is known to specialize on nutrient poor pine forests have low densities of other birds (Lehikoinen et al. 2017).

**Figure 2.**
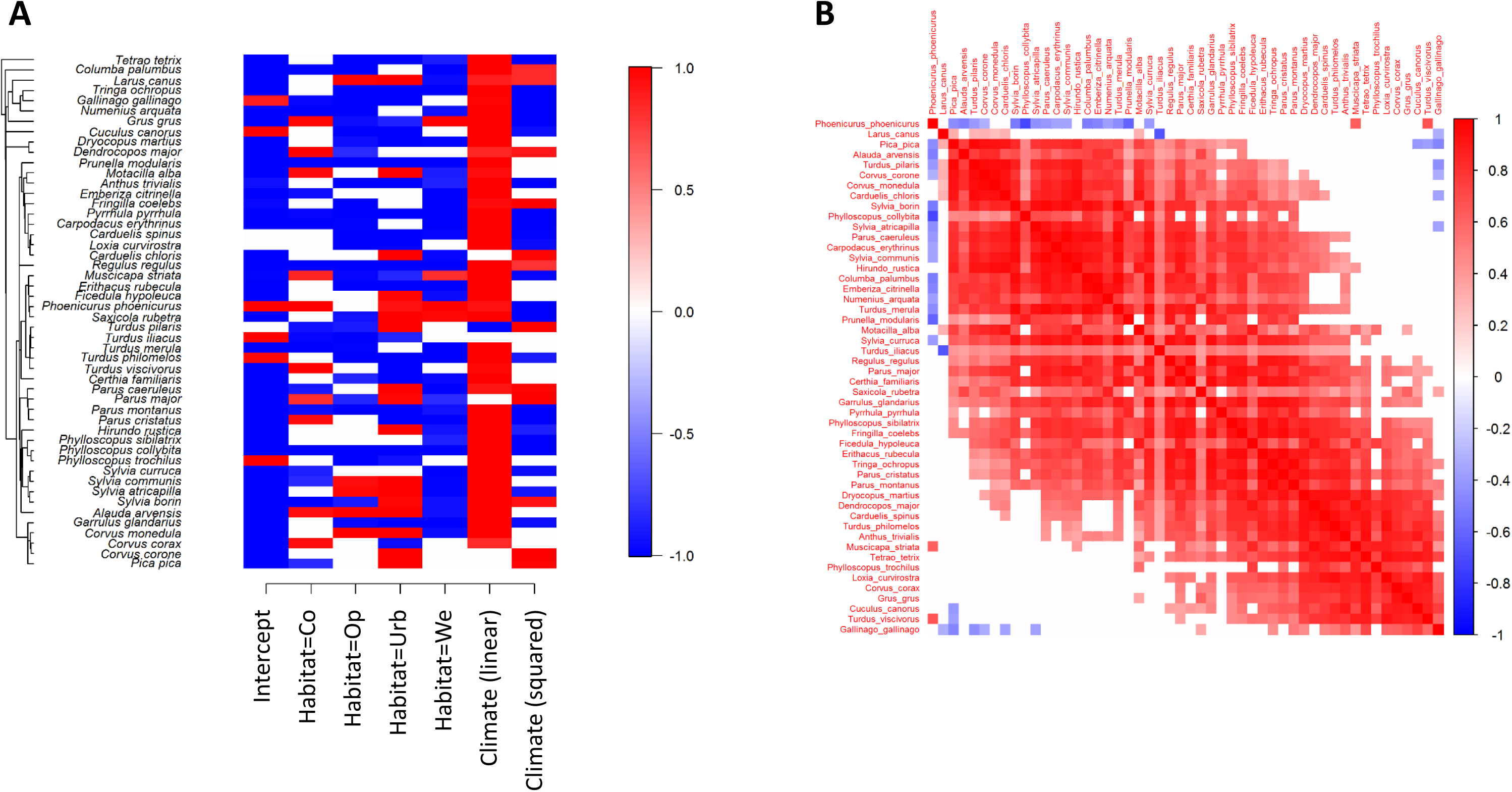
Exploring parameter estimates. Panel A shows those *β*-parameters (species responses to environmental covariates) that received in model PA.X a high (at least 95% posterior probability) statistical support of being positive (red) or negative (blue). The species are ordered according to their phylogeny shown on left. Panel B illustrates the residual association structure in model PA.S, with positive associations with high (at least 95% posterior probability) statistical support shown by red and negative associations by blue. The species have been ordered in a way that best illustrates the association structure of the data.

### Step 5. Making predictions

HMSC-R includes functions that make and illustrate predictions over categorical (here the habitat type; Fig. 3ABCD) as well as continuous (here spring temperature, results shown in the vignette) environmental gradients. Concerning the spatial predictions (Fig. 3EFGH), we have used here a model fitted to data from 200 locations to predict species occurrences in ca. 10,000 locations. As we have generated these predictions with model PA.XS, they utilize both environmental and spatial information. Predictions can be illustrated at the level of individual species, e.g. showing that *Corvus monedula* prefers urban habitats (Fig. 3A) and mainly occurs in Southern Finland (Fig. 3E). Predictions can also be illustrated at the level of species richness, showing that urban habitats host the most species and wetlands and open habitats the least species (Fig. 3B), and a decreasing gradient from South to North (Fig. 3F). Predictions can further be illustrated at the level of community-weighted mean trait values, showing e.g. that the proportion of resident species is lowest in wetlands (Fig. 3C) and decreases towards the North (Fig. 3G). One way to visualize variation in community composition is to cluster the predicted communities into a discrete set of communities of common profile. Doing so shows that urban areas have distinct communities (Fig. 3D), and that communities are structured along the latitudinal gradient in terms of their species composition (Fig. 3H).

**Figure 3.**
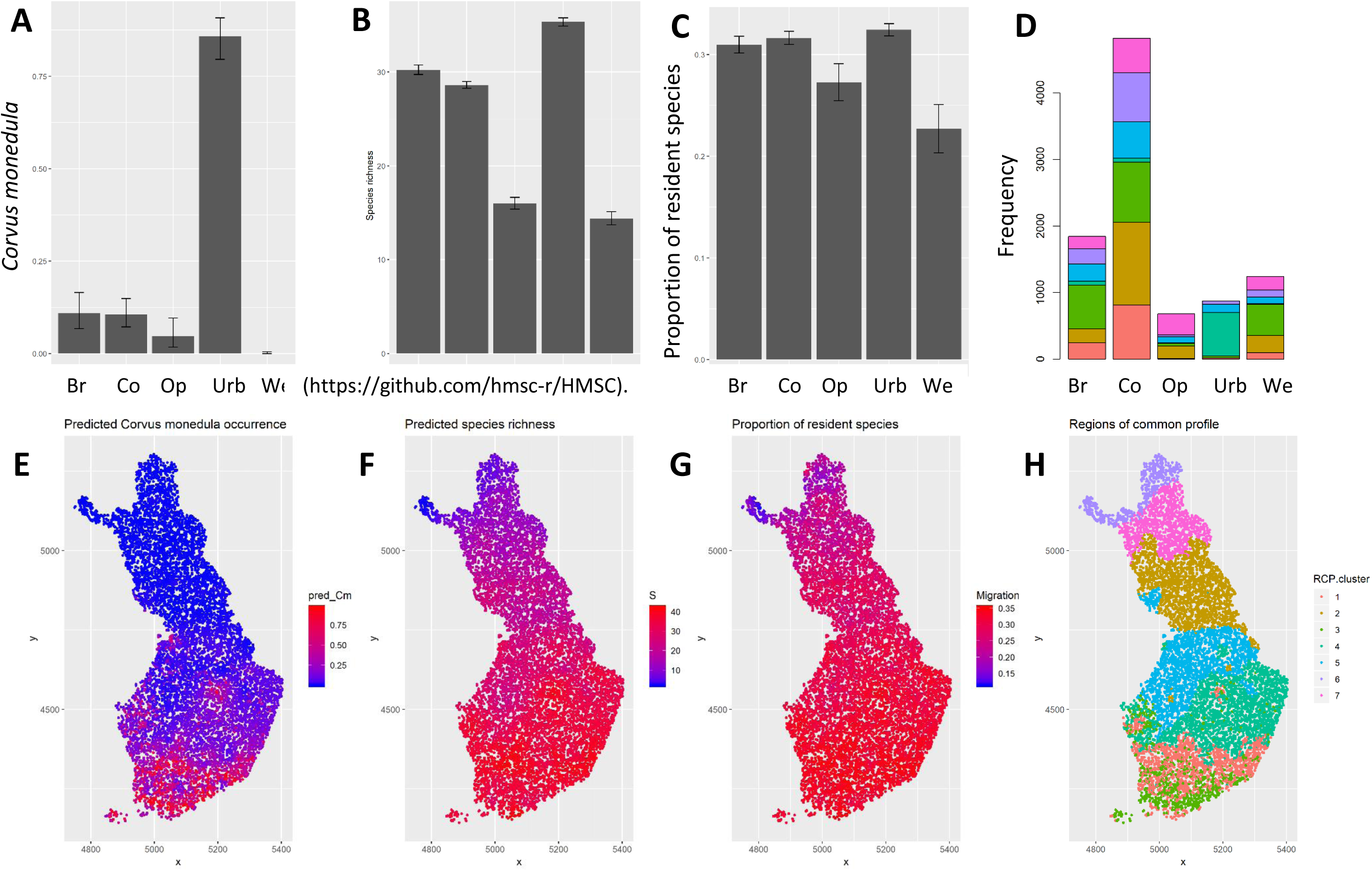
Making predictions. The upper panels exemplify predictions over environmental gradients, and lower panels spatial predictions, both of which can be used to quantify the influence of covariates on species occurrence (AE), species richness (BF), community weighted mean traits (CG), or regions of common profile (DH). The scenario simulations are used here to predict species communities in different habitat types, whereas the spatial predictions are used to predict species communities on a grid covering entire Finland (the training data come from a small subset of 200 locations). The panels ABC and based on Model PA.X and the remaining panels on Model PA.XS.

## Conclusion

At present, the most widely applied methods for data analysis in community ecology are based on ordination approaches. While we appreciate the value of ordinations as a fast way for making descriptive analyses, we encourage community ecologists to make even more out of their data by applying also model-based approaches, especially the newly emerging JSDMs (Warton et al. 2015). We hope that the users find HMSC-R 3.0 to provide a highly functional and user-friendly package for doing so.

## Author Contributions

GT designed and implemented the main part of the HMSC-R 3.0 software. OO contributed specific functions to HMSC-R, implemented the first versions of the simulated and real data case studies and devised the first drafts of the simulated vignettes. ØO devised the first draft of the bird vignette, revised all vignettes, and contributed specific functions to HMSC-R. NA and OO wrote the first version of the manuscript. AL contributed the data for the case study. All authors contributed significantly to the writing of the final version of the manuscript.

## Acknowledgements

This work was funded by Academy of Finland (grants 284601 and 309581 to OO and 275606 to AL), Jane and Aatos Erkko Foundation (grant to OO), and the Research Council of Norway through its Centres of Excellence Funding Scheme (223257) to OO via Centre for Biodiversity Dynamics. Finnish Ministry of Environment has supported the census scheme.

## Data accessibility

HMSC-R 3.0 is available at Github (https://github.com/hmsc-r/HMSC). The vignettes 1-5 are also placed there.

